# Social Perspectives on Larvicide-Based Mosquito Control in Urban Quebec

**DOI:** 10.1101/2025.10.20.683413

**Authors:** Francesca Sotelo, Ann Lévesque, Megan Larouche, Jérôme Dupras, Katrine Turgeon

## Abstract

The spreading of *Bacillus thuringiensis israelensis* (Bti) in urban waterways, as a biological larvicide to control biting insect populations and mitigate disease transmission, has been subject to scientific scrutiny due to its potential environmental impacts. Public and political discussions have revealed recurring opinions that reflect broader social perspectives. While scientific research on Bti progresses, knowledge of public understanding and perspectives, which are important for decision-making, remains limited. The main objective of this research was to identify and compare the social perspectives of Quebec citizens, primarily in the city of Gatineau, regarding the use of Bti for mosquito and black fly control in aquatic environments. We used Q methodology, a mixed approach combining both quantitative and qualitative methods to identify social perspectives related to the use of Bti. Our analysis revealed three social perspectives, which account for 51% of the explained variance. They reflect the broader range of opinion within citizens, and are described as “The critical environmentalist”, “The diligent mosquito hunter”, and “The Bti enthusiast”. Each accounting for 29%, 12% and 10% of the explained variance, respectively. The common thread across all three perspectives is the minimal public awareness and understanding of Bti and its environmental implications. Providing a first characterization of social perspectives in North America regarding larvicides, this research contributes to a more comprehensive understanding of public opinion and informs the development of inclusive and transparent policies.

## 1. Introduction

By increasing temperatures and altering precipitation regimes, climate change is responsible for the increase in biting insect density, notably mosquitoes, and the expansion of their ecological niches worldwide (Field et al., 2019). Increased temperatures are promoting ideal conditions for mosquito breeding and survival, with optimal temperatures ranging from 20 to 27°C, depending on species (Couper et al., 2024; World Health Organization, 2023). An increase in mosquito densities leads to an increased incidence of vector-borne diseases such as malaria, filariasis, dengue fever, yellow fever, Zika virus, Chikungunya, and West Nile virus (Valtierra-de-Luis et al., 2020; Ryan et al., 2019). Out of the roughly 400 known mosquito species, about 30 are capable of spreading diseases to humans (Valtierra-de-Luis et al., 2020). The primary vector genera include *Anopheles*, *Culex*, and *Aedes*, which belong to the Anophelinae and Culicinae subfamilies (Valtierra-de-Luis et al., 2020).

The increase in mosquito density has disproportionate negative effects in tropical and subtropical climates compared to temperate and boreal ones. In the tropics, according to the latest report from the World Health Organization (WHO), malaria cases in 2023 were estimated at 263 million malaria cases and 597,000 deaths worldwide (World Health Organization, 2024). Approximately 94% of malaria cases and 95% of deaths were recorded in Africa (World Health Organization, 2024). In temperate and boreal climates, the increase in *Aedes* density has a much less dramatic consequence on humans. Mosquitoes are mostly linked to nuisance for human outdoor activities, as well as human discomfort and irritation (Halasa et al., 2014). Their seasonal outbreaks can impact tourism, outdoor labour for seasonal workers, and overall appreciation of wetlands (Brühl et al., 2020).

The adoption of insecticides to control biting insect populations was met with reluctance, mostly due to the devastating environmental effects of DDT, which came to light in the 1950s and led to increased public and regulatory caution (Dambach et al., 2018). However, driven by public health concerns, there has been a global increase in the deployment of biological larvicides and insecticides. *Bacillus thuringiensis israelensis*, commonly referred to as Bti, is a bacterium discovered in Israel in 1976. As of the early 1980s, it has been commercially available and used worldwide as a biological larvicide to control biting insect populations, specifically targeting the Diptera family (Lacey & Merritt, 2003; Goldberg & Margalit, 1977). Bti is sprayed in waterways and wetlands where biting insects, such as mosquitoes and black flies, complete their early stages of development (larvae). Once ingested by the insect larvae, the bacterium’s crystallized proteins dissolve in the gut, releasing toxins that cause fatal damage to cell membranes (Valtierra-de-Luis et al., 2020; Boisvert & Lacoursière, 2004). These proteins can affect multiple insect species belonging to different orders, like Lepidoptera, Diptera, Coleoptera, Hymenoptera, Hemiptera, Orthoptera, and other organisms such as mites and nematodes (Valtierra-de-Luis et al., 2020).

Although Bti is reported to have reduced effects and low toxicity on non-targeted animals (Després et al., 2011; Goldberg & Margalit, 1977), its ecological effects remain controversial. There is cumulative empirical evidence that Bti and its adjuvant ingredients can affect the whole food web depending on species sensitivity and experimental conditions, especially when applied at higher concentrations or with specific formulations (Klein & Cabrera, 2023). Biting insects are an important food source for both aquatic and terrestrial animals, and some species play a significant ecological role as pollinators in their adult stage (John et al., 2024; Shaalan & Canyon, 2009). Bti also eliminates important non-biting insect prey, such as chironomids that belong to the Diptera family, and this can indirectly impact the demography of their predators, such as Odonata (Brühl et al., 2020; Allgeier et al., 2019; Jakob & Poulin, 2016) and insectivorous bird species (Poulin et al., 2010). Bti may also pose toxicity risks to amphibians (Allgeier et al., 2018; Lajmanovich et al., 2014). Finally, another concern with the use of larvicides is the increased resistance among biting insect populations. Continuous use of the same bacteria can lead to the selection of treatment-resistant individuals, which eventually results in resistant and stronger *Aedes* populations (Carvalho et al., 2025; Brühl et al., 2020).

The ingredients in Bti products (e.g., formulants and adjuvants) are kept confidential (Health Canada, 2011). Ethically, this form of protection, whether through trade secrets or patents, raises concerns about the transparency of its actual effects on the environment and has led to suspicions in the scientific community. The choice to use or not use Bti in this instance is not merely a technical matter, but it might be one of competing values and interests. Ravetz (2004) refers to this kind of situation as one of high decision stakes, in which particular actors (i.e., companies, advocacy organizations, or government agencies) have a great deal to gain or lose. In such cases, those with higher gains or losses could emphasize uncertainty about the product or soften particular risks as a rationale for their position.

When decision stakes are high, it is strongly suggested to extend the knowledge process beyond the scientific community and draw on locally based expertise and diverse perspectives, including those of the general public (Lidskog & Berg, 2022; Ravetz, 2004). Although the inclusion of the general public does not necessarily complement scientific knowledge, taking the pulse of the public provides important insights into their concerns and interrogations (Marres, 2007). This, in turn, ensures that public opinion is considered, and that proposed solutions are relevant, legitimate, understood by the population (Lidskog & Berg, 2022).

The objective of this project was to identify the social perspectives related to the use of Bti for mosquito and black fly control in wetland and water environments in Quebec, Canada (temperate climate). We aimed to answer two main questions: 1) What are the different social perspectives related to the use of Bti for mosquito and black fly control in wetlands and water environments in Quebec? and 2) Are there areas of convergence and divergence among the general public regarding the use of this bioinsecticide? Answering these questions will enhance our understanding of public opinions regarding larvicide application, providing insight to inform environmental decision-making.

## 2. Materials and methods

The general public’s perspectives can be captured in various ways. These could include direct interaction (focus groups or interviews), analysis of media coverage, frequently asked questions, or comments on forums or social media, as well as surveys and questionnaires (Maranta et al., 2003). On the one hand, qualitative approaches, such as one-to-one interactions, can be time-consuming, and quantitative approaches, like surveys, provide few clues as to how the general public’s opinions are formed. With the Q methodology, this research used a mixed-methods approach combining both quantitative and qualitative methods to identify social perspectives related to the use of Bti. This method, developed by the physicist and psychologist William Stephenson (Stephenson, 1935), is particularly useful for exploring social phenomena, as it enables the characterization of discourses within a population on a given issue and identifies the points of consensus and divergence between each of the identified discourses. Q methodology is employed in various scientific disciplines, including political science, environmental science, and social science. Q methodology bridges the gap between qualitative and quantitative research.

The qualitative component of the Q methodology enables participants to express their subjective point of view, whereas the quantitative component provides insight into how opinions are formed and expressed. This method uses statistical techniques, such as correlations and factor analysis, to support Q methodologists in identifying the main discourses present within a group of participants and to examine the variation between them (Watts et al., 2012). This said, a case study approach and a mixed methodology were used to conduct this research. Data collection was confined to a specific territory of the province, namely the city of Gatineau, Quebec’s fourth-largest city.

### 2.1. Case study: The controversial use of Bti in Gatineau, Quebec, Canada

In northern countries like Canada, the risk of deadly diseases transmitted by mosquitoes is minimal. However, the West Nile virus has remained a concern since its first appearance in 2002 and is now present in multiple regions in the province of Quebec (Gouvernement du Québec, 2023). In most cases (80%) in Quebec, the infected person has no symptoms (Gouvernement du Québec, 2023). Less than 1% of infected people will develop an illness, such as meningitis or encephalitis, with significant neurological effects (Gouvernement du Québec, 2023). Even so, Bti has been used for decades in streams of multiple Quebec municipalities since 1982 (Boisvert & Lacoursière, 2004). The primary reason is to reduce the nuisances caused by biting insects, thereby improving the comfort of citizens and tourists, which in turn helps the economy. However, in the current context of a global biodiversity loss (IPBES, 2019), the eradication of biting insect populations is highly disputable, and the use of Bti is being reconsidered, especially since rich biodiversity inhabits streams and wetlands.

In Canada, pesticides are rigorously regulated to ensure that the risks they pose to human health, and the environment are minimal. The Pest Management Regulatory Agency (PMRA), a division of Health Canada, oversees all aspects of pesticide regulation, including the evaluation, registration, and re-evaluation of pest control products, in respect of the *Canadian Environmental Protection Act* (CEPA; S.C., c. 33; Health Canada, 2011). The PMRA conducts science-based risk assessments to determine whether a pesticide meets safety and efficacy standards before it can be approved for use (Health Canada, 2011). This includes a rigorous, science-based evaluation of the product’s formulation provided by manufacturers. However, in the case of Bti, information on the formulant is classified as a trade secret and the disclosure of this type of information to the general public is prohibited under the *Access to Information* (ATIA; R.S.C., c. A-1) *and Privacy Act* (R.S.C., c. P-21; Health Canada, 2011). To maintain safety standards, the PMRA re-evaluates approved pesticides every 15 years, ensuring compliance with the most current regulations and scientific findings (Health Canada, 2011). Most formulations of *Bacillus thuringiensis israelensis* (Bti) used for the control of mosquito and black fly larvae are classified as “restricted” in Canada and must be applied directly to aquatic habitats where larvae are present (Health Canada, 2011). In many provinces, individuals applying restricted-use pesticides must hold certification, and in some cases, a permit from the provincial pesticide regulatory authority.

In the province of Quebec, the use of Bti is regulated by two laws, which are the respect of the *Environmental Quality Act* (EQA; RLRQ, c. Q-2) and the *Act respecting the conservation and development of wildlife* (RLRQ, c. C-61.1). The reinforcement of these laws is mandated by the Ministry of the Environment, the Fight against Climate Change, Wildlife and Parks (MELCCFP). The MELCCFP’s regional wildlife management branches must issue authorizations or wildlife advisories when projects involving the control of biting insect populations are proposed (Klein & Cabrera, 2023). Quebec guidelines for controlling biting insects with Bti requires a case-by-case approach to balance public health, human well-being, and environmental protection (MELCCFP, 2023). These guidelines specify the person or organization requesting authorization need to seek for social acceptability of their Bti use. According to the government of Quebec, social acceptability is defined as “the outcome of a collective judgment or collective opinion of a project, plan or policy” (Gouvernement du Québec, 2025). While social acceptability is not quantifiable and can only be described, it remains a decisive factor in whether a project, plan, or policy is accepted or rejected. Moreover, the precautionary principle (*i.e.,* to implement measures that avoid potential and irreversible threats to the environment in the future; CEST, 2022) is increasingly adopted and promoted, putting a hold on the use of Bti in multiple cities. This is the case in the city of Gatineau, located in southern Quebec.

In the Gatineau region, Bti has primarily been used in the sectors of Masson-Angers and Gatineau, covering seven electoral districts (Pointe-Gatineau, Carrefour de l’Hôpital, du Versant, Bellevue, Lac Beauchamp, la Rivière-Blanche, and Masson-Angers), as shown in Figure 1 (Ville de Gatineau, 2023). The spreading of Bti has occurred every summer from 1996 up to 2023 (Ville de Gatineau, 2023). The expenses were mostly covered by a mosquito tax paid by the landowners living in these districts (Ville de Gatineau, 2023). During those years, Bti was promoted as a safe product that effectively eradicated mosquitoes to improve human comfort, while exerting no significant risks to the environment or human health. However, accumulating evidence from scientific research has brought many doubts regarding the lack of effects on the environment, leading to a rethinking of its usage (Klein & Cabrera, 2023; Duguma et al., 2015).

**Figure 1.**
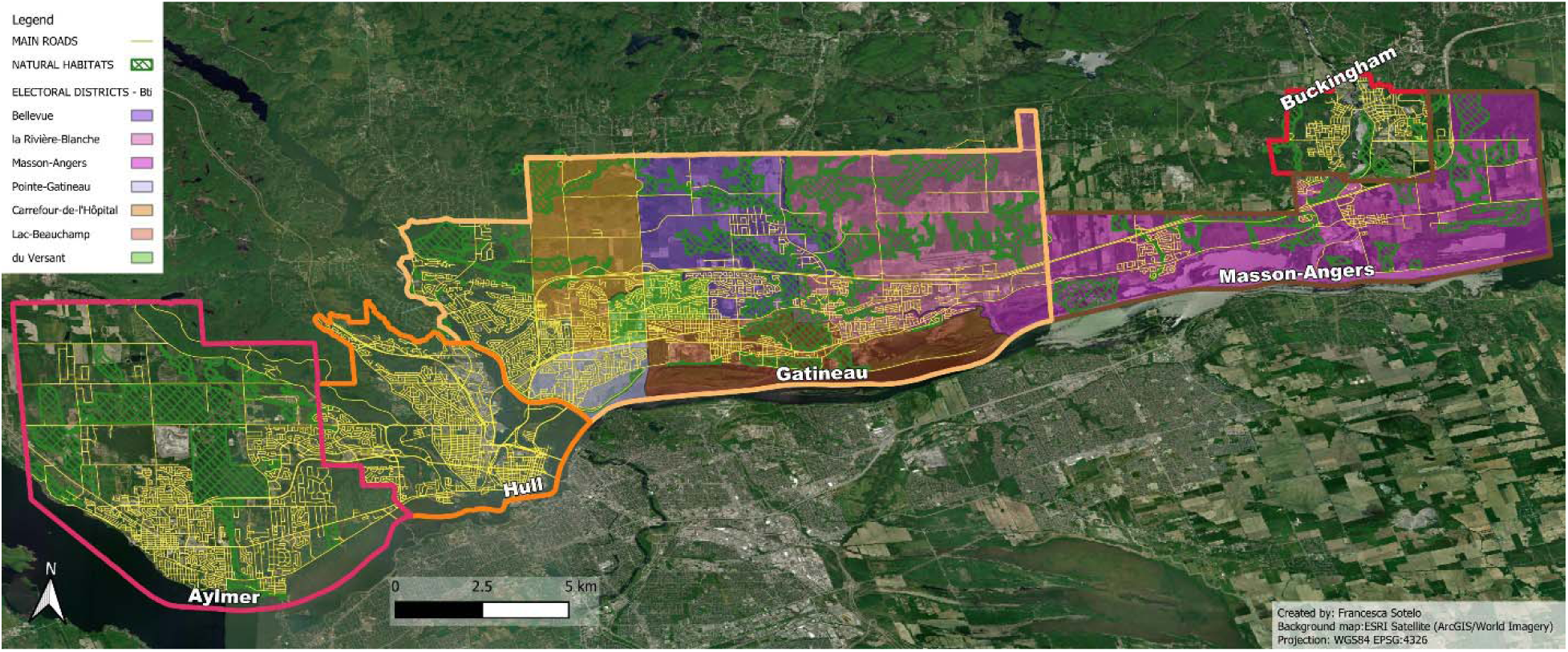
Map of Gatineau divided into its sectors and electoral districts where Bti was spread.

As of April 2023, following a public consultation in 2022 on Gatineau’s biodiversity plan for 2024-2028, which included an assessment of the social acceptability for pesticides, the municipal council of Gatineau has decided to put an end to the biting insect biological control program, thereby prohibiting the use of Bti in its districts (Ville de Gatineau, 2023). The cessation of Bti usage in Gatineau was based on a precautionary principle, therefore due to a lack of certainty, excessive ambiguity, and conflicting research regarding its impacts on the food web. Since then, the council has been exploring other options to reduce the discomfort caused by mosquitoes, such as installing fly traps, and concentrating their efforts in the districts that are most affected.

The public consultation allowed a defining conversation for Gatineau’s biodiversity action plan (Ville de Gatineau, 2024). The first phase of the public consultation consisted of an online group discussion held during a video conference on October 12th and 13th, 2022. These discussions were to gather ideas, general preoccupations, and the main priorities for Gatineau’s biodiversity action plan from different groups. Twenty-eight people participated in these discussions, comprising environmental groups, city partners, and citizen and resident associations. The second phase consisted of an online survey. This online public consultation, conducted in Gatineau between August 26, 2022, and October 31, 2022, received over 300 responses to the questionnaire, almost 80 submissions of ideas, and five essays reflecting on the biodiversity action plan (Ville de Gatineau, 2024). This survey demonstrated that most respondents (over 98%) believed biodiversity is important, and in both consultations, they agreed that the use of pesticides in urban areas should be better regulated (Ville de Gatineau, 2024).

Following this public consultation, thirteen objectives for Gatineau’s biodiversity plan were established, such as to protect 30% of Gatineau’s terrestrial and aquatic environments, which aligns with the international target of the Biodiversity Framework adopted during COP 15, and to restore 30% of degraded environments, while also compensating for destroyed environments (Service de la transition écologique & Nature Québec, 2024; Ainsworth et al., 2022). Another objective, strongly related to the use of Bti, is to reduce sources of pollutants that threaten the integrity of ecosystems, and doing so by reducing pesticide use. Measures that will be implemented include the adoption and enforcement of a pesticide bylaw, as well as informing citizens about alternatives to pesticides (Service de la transition écologique & Nature Québec, 2024).

The choice to limit this case study to Gatineau, with few respondents living in rural areas within the Outaouais region, was justified by this important ongoing and topical issue, as well as the necessity of gathering and understanding social perspectives from the general public to inform decisions regarding their biodiversity conservation plan.

### 2.2. Mix-method steps

The research was conducted in three steps between January and April 2023 (Table 1) to generate relevant and meaningful statements in the Q-method.

**Table 1.**
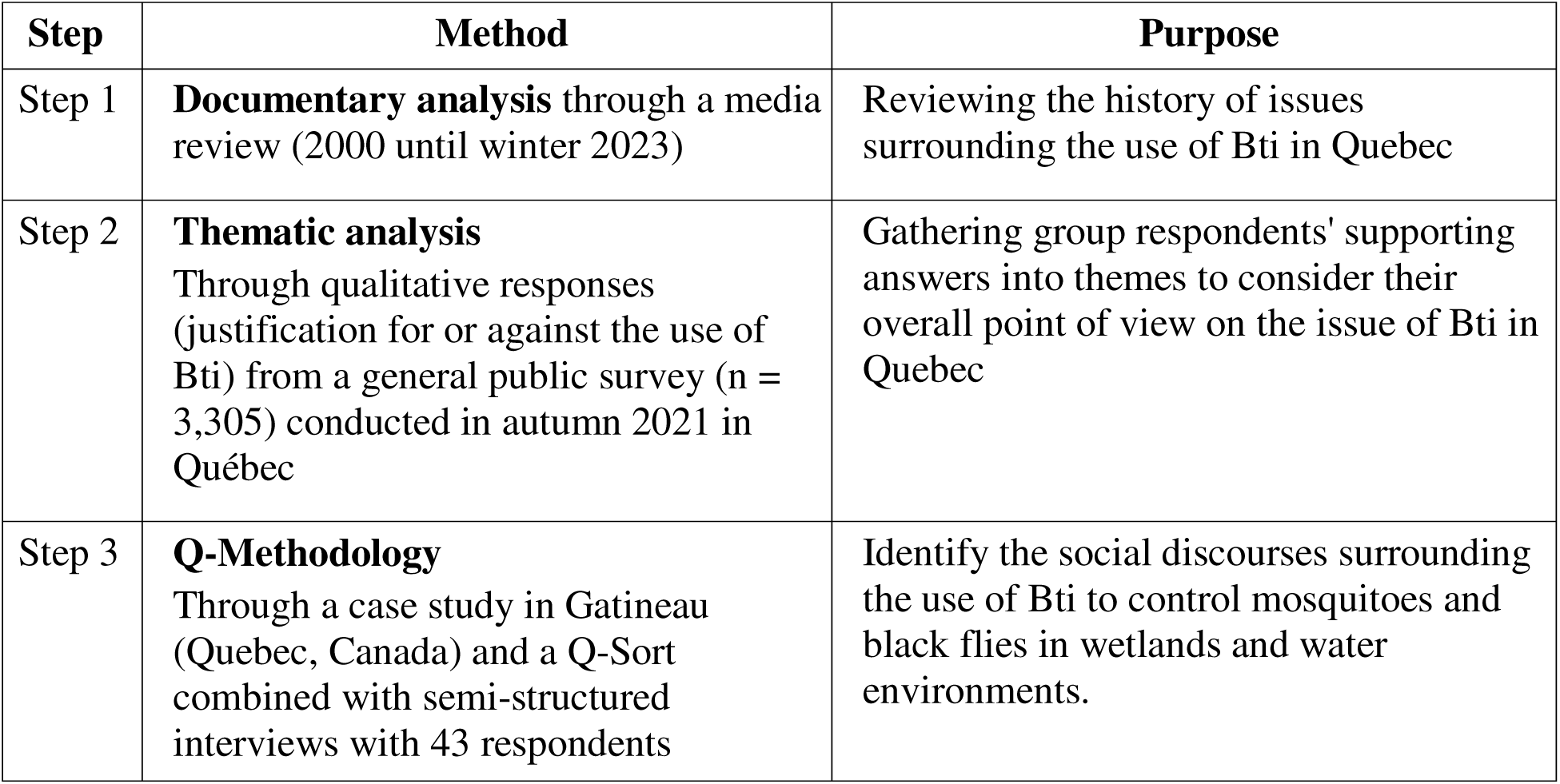
Mix-method steps summarized.

#### 2.2.1. Step 1: Documentary analysis

The first step involved a review of Quebec media on the use of Bti, covering the period from 2000 to 2023. Nearly 50 newspaper articles were analyzed to familiarize researchers with the various issues surrounding Bti in Quebec. This process was carried out using Eureka software, a digital news database developed by “Bibliothèque et Archives Nationales du Québec”.

#### 2.2.2. Step 2: Thematic analysis

The second stage involved conducting a thematic analysis to provide an exhaustive portrait of the various opinions on the use of Bti among the Quebec population. To conduct this analysis, we utilized qualitative responses (justifications for or against the use of Bti) from a survey on wetlands conducted in the fall of 2021, targeted to the general public (Canada Research Chair in Ecological Economics). A total of 3,305 respondents took the online survey in October 2021. While the survey was not specifically targeted at Bti, its main purpose was to assess public support in Quebec for wetland conservation at the regional level, including expansion, willingness to pay, and preferences for wetland conservation and restoration actions beyond current government action (Gagné, et al., 2022). The questionnaire contained a specific part on Bti, in which background information was given to the respondents at first to better understand the issue:

> *"To reduce the number of nuisance insects (black flies and mosquitoes) near dwelling areas, certain municipalities in Quebec spray ponds, lakes, rivers, and marshes with Bti, an insecticide whose action kills the larvae of these insects. However, this use has broader environmental implications, since many species depend on these insect populations for survival."*

After reading this brief background, the respondents were asked to state to what degree they agreed with the use of Bti in wetlands to eliminate black flies and mosquitoes. Three response options (yes, no) were provided, and respondents were invited, if they wished, to provide the reasons for their answer via an open-ended text box. Respondents also had the choice to select the answer “I have no opinion on this matter” without leaving any reasons.

Out of the 3,505 respondents from all regions of Quebec, 702 were in favour of using Bti, 1,680 were against it, and 1,123 were undecided. Out of those who did take a position, 454 respondents in favour provided at least one reason to justify their choice, while 1,140 respondents opposed also left a comment explaining their position. Based on these qualitative comments, a thematic analysis was carried out. The development of themes was carried out via NVivo in pairs across five teams. Each team member employed a continuous thematization technique, which involves reading each respondent’s justification answers and creating themes as they progressed through their analysis (Paillé & Mucchielli, 2016). A triangulation of the themes that emerged was carried out by each team to increase the robustness of this step. Ultimately, the principal investigator of this research consolidated the themes identified by the five teams and validated them collectively. In total, 28 themes were identified (Appendix A), representing all of those found during the group thematization exercise.

#### 2.2.3. Step 3: Q-Methodology

Q methodology has gained prominence in research addressing complex and disputed socio-ecological issues, offering a structured and statistical approach to exploring varied viewpoints (D’Amato et al., 2019). Central to this methodology is the Q-set technique, wherein participants rank a set of predefined statements along a continuum of agreement (typically from +4 to −4) in the form of a pyramid (Appendix B), which forms a Q-sort when completed (Brown, 1996). The statements were developed prior to the interviews, from the thematic analysis detailed above. The Q-set has been designed to be representative, balanced, and comprehensive. The representative component of a Q-Set requires that all statements provide good coverage of the research question under study, while the balanced component requires that the full range of possible opinions be present in the Q-Set as well (Watts & Stenner, 2012). The Q-sorts were then subject to factor analysis to identify shared viewpoints, or "archetypes," based on common patterns in the placement of statements, allowing the calculation of correlations between individuals rather than between variables.

Before classifying the statements, respondents were given a context for the project. They were informed that Bti is a bio-insecticide used to control mosquitoes and black flies. It was specified that this insecticide is biological, as its active ingredient is a bacterium naturally found in the soil, but this bacterium kills the larvae of these insects, preventing them from reaching the adult stage. Respondents were also informed that for several years, many Quebec municipalities have been spraying ponds, lakes, rivers, and wetlands with Bti to control populations of these biting insects. Respondents were also advised that the use of bio-insecticides to control mosquitoes and black flies in wetlands and waterways is not widely accepted by the Quebec population. In support of this statement, respondents were informed that a survey of 3,505 citizens was conducted in 2021 to determine the reasons justifying the application or non-application of Bti, and that the statements to be ranked during the interview summarized the overall justifications of the Quebec population gathered from this survey.

Before ranking, respondents were informed that there are no right or wrong answers in the ranking process. First, respondents were asked to classify each statement into one of 3 categories: agree, neutral, or disagree. Respondents were then invited to reread each statement and classify it in an appreciation grid (Appendix B). This grid contains 39 boxes, i.e., the total number of statements to be classified, and an appreciation scale divided into 9 preference classes (−4 = most disagree, 0 = no opinion, + 4 = most agree). After sorting, respondents were asked to rank the statements at the extremes (+3, +4 and −3, −4) of the sorting grid, to list and explain the two statements that surprised them most, and to validate whether the Q-Set seemed complete.

### 2.3. Statistical Analyses

All statistical processing was performed using PQMethod version 2.35 (Schmolck, 2014). Analyses were carried out in five steps. First, each Q-sort was correlated with the set of collected Q-sorts to generate a correlation matrix (Table 3; Watts & Stenner, 2012). Second, principal component analysis (PCA) was conducted to determine axes explaining the most variation in the data, based on the Kaiser-Guttman criterion (eigenvalues > 1). Third, three factors were retained using Horne’s parallel analysis and the researchers’ judgment, based on their experience with each respondent during the interviews (Watts & Stenner, 2012). Fourth, principal component analysis was performed to find the strongest correlations among the different Q-sorts. To determine the location of each respondent for the 3 selected factors, Humphreys’ rule (a significant score ≥ (2.58 / √(□□□ of statements): 0.37) was used. Fifth, we performed a varimax rotation to increase possible correlations (Watts & Stenner, 2012) and maximize the correlation of individual points of view similar to one another (Brown, 1993; Davies, 2017). Detailed information on statistical principles specific to the Q-method can be found in Brown (1980).

## 3. Results

Data were collected from 59 respondents over a four-week period between March and April 2023. The majority of participants resided in Gatineau, with only six respondents living in rural areas outside the city but still within the same region. All interviews with respondents, during which Q-sorts were completed, were conducted by eleven university students as part of their curriculum. The method was standardized across all students, using the same forms and introductory scripts at the beginning of each interview to ensure comparable data. Respondent profiles corresponding factor scores are presented in Appendix C. The results of the PCA revealed three respondents that did not correspond to a specific profile and overall three social perspectives that reflect the broader range of opinion, which have been titled as “The Critical Environmentalist” (factor 1), “The Diligent Mosquito Hunter” (factor 2), and “The Bti Enthusiast” (factor 3). The highest-ranked statements for each perspective are presented in Table 2. It is important to note that a perspective is described not only by the statements ranked at the extremes but also by the entire ranking, including those placed in the neutral zone. The Factor z-score of all statements with their respective ranking can be found in Appendix D. These results explain 51% of the variance, which is considered satisfactory for Q-methodology (≥ 35%; Watts & Stenner, 2012). However, the remaining 49% reflects additional opinions and perspectives not captured by the three main factors or perspectives. These viewpoints are not insignificant or less important but rather represent less commonly shared opinions or those not highlighted by the selected statements or analyses. The following sections present the three social perspectives that arise from this analysis.

**Table 2.**
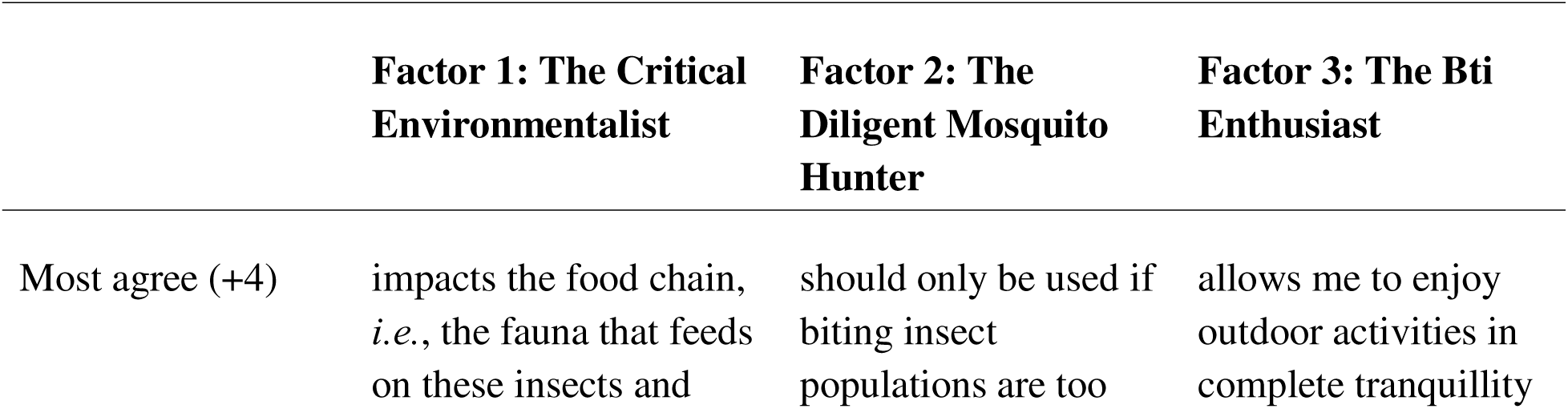
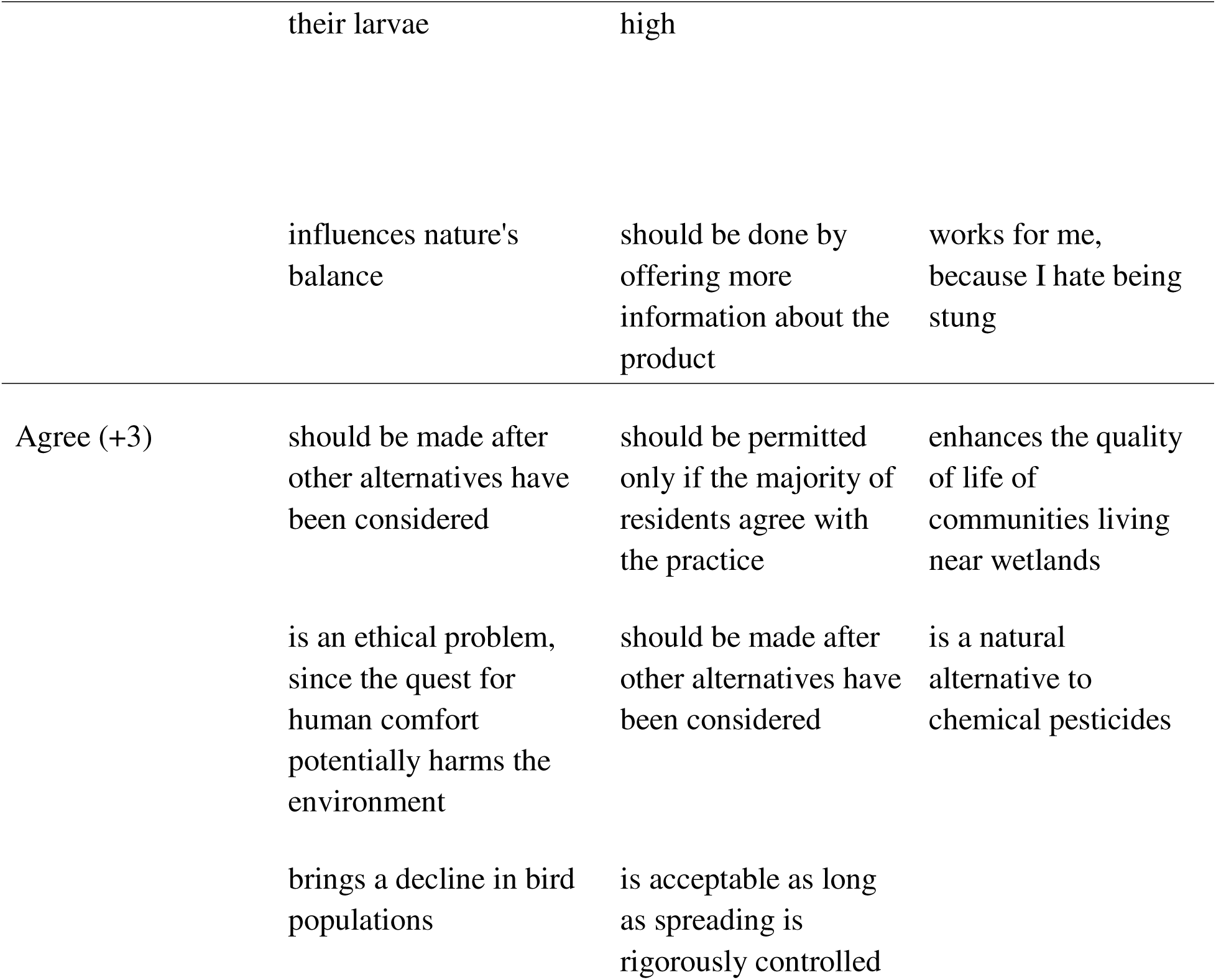
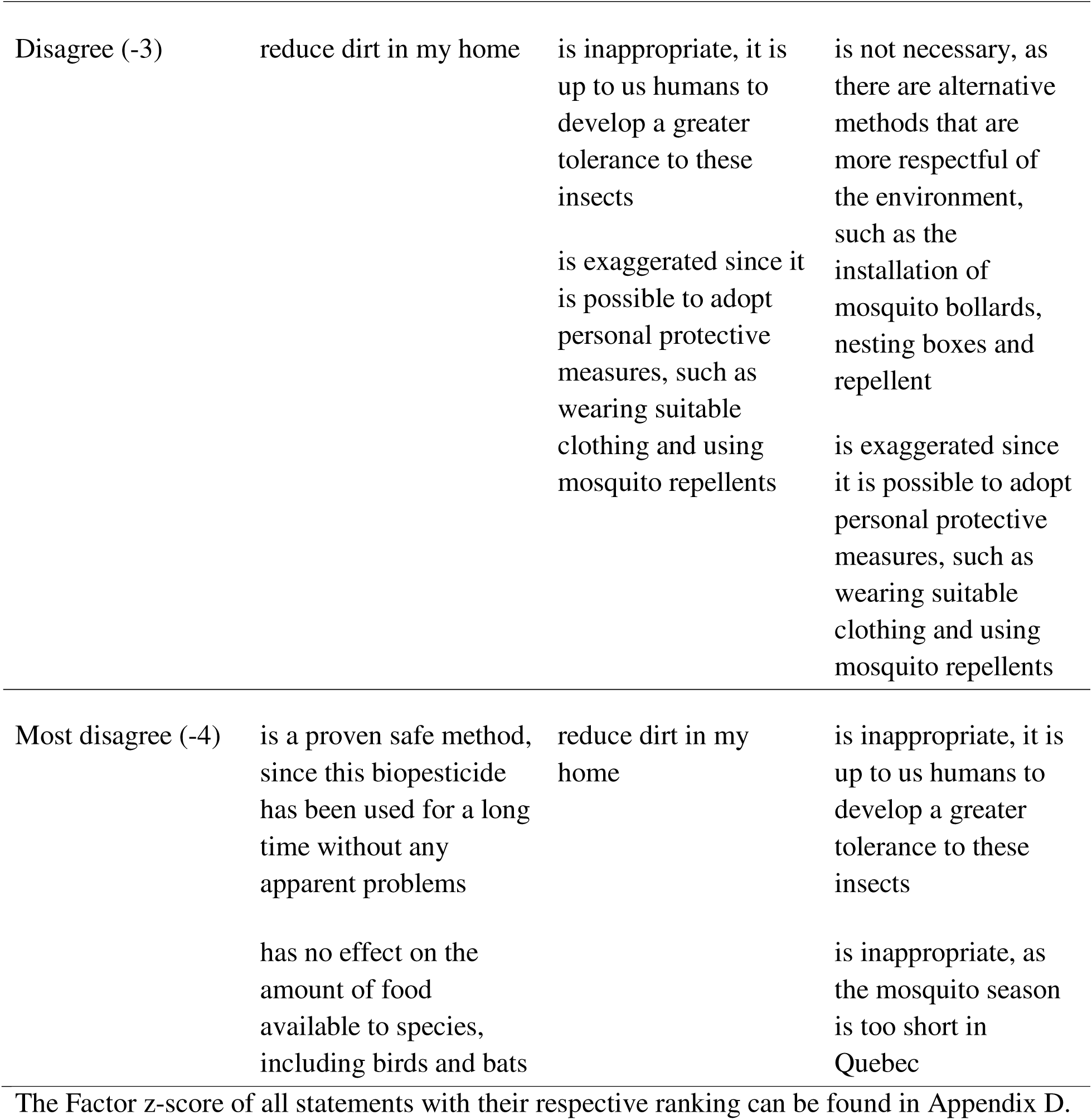
Highest-ranked statements for each perspective.

### 3.1. Social perspective 1: The Critical Environmentalist

The first perspective accounts for 29% of the explained variance and includes 36 defining variables, *i.e.*, the number of respondent Q-Sorts corresponding to this viewpoint (Appendix C). It clearly expresses that the use of Bti raises ethical concerns, as it compromises wildlife for the sake of human comfort and needs. Respondents aligned with this viewpoint remain neutral regarding the extent to which Bti improves the well-being of people engaged in outdoor activities or living near wetlands during mosquito season. They strongly support the use of alternative methods with fewer environmental impacts and express skepticism toward Bti’s safety, regulation, and current applications. According to this perspective, citizens should be better informed about the product, and its use should be considered only as a last resort. While acknowledging Bti’s potential in controlling mosquito-borne diseases and protecting human health, this perspective places greater emphasis on its ecological consequences, particularly its effects on food webs, pollinators, and how it undermines ethical consideration for animal life. Given these critical and environmentally conscious beliefs, this perspective has been labeled *the critical environmentalist*.

### 3.2. Social perspective 2: The Diligent Mosquito Hunter

The second perspective accounts for 12% of the explained variance and includes 11 defining variables (Appendix C). This perspective believes that the use of Bti is necessary when the population of biting insects is too abundant, provided that affected citizens are consulted beforehand. This perspective agrees that treating areas with high concentrations of mosquitoes and black flies allows residents to enjoy nature more fully. While Bti is considered a preferable alternative to chemical pesticides, it is not necessarily perceived as safer. Respondents belonging to this perspective favour environmentally friendly and health-conscious alternatives, although they are less inclined toward traditional methods, such as protective clothing or insect repellents. Instead, they express interest in options such as mosquito repellent plants, nesting boxes, and bollards. This group supports the use of Bti when no other viable solution is available but remains relatively unconcerned about its potential effects on wildlife. Nonetheless, access to clear and detailed information before application is considered essential. Valuing transparency and public consent, this perspective underscores the importance of informing and involving citizens. Reflecting these principles, it is referred to as *the diligent mosquito hunter*.

### 3.3. Social perspective 3: The Bti Enthusiast

The third perspective accounts for 10% of the explained variance and includes nine defining variables (Appendix C). This perspective is composed of individuals who express a strong aversion to mosquitoes and black flies, viewing a life without biting insects as offering significant individual and collective benefits. They recognize its potential to enhance the image of wetlands and improve the well-being of farm animals. All other conventional alternatives proposed for repelling biting insects are not suitable for this perspective. Within this group, Bti is perceived as a natural product, though respondents remain neutral regarding its regulations and the safety associated with its use. Ethical concerns about Bti are largely absent as long as it effectively reduces insect populations. However, they remain neutral about its efficacy, which might be explained by the lack of knowledge regarding this product. Participants who align with this view do not actively seek additional information about Bti and do not believe its use should be restricted due to limited scientific knowledge. While they acknowledge potential ecological impacts, such as reduced food availability for wildlife, and possible effects on pollinators and bird populations, they slightly disagree that Bti significantly harms human health, the environment, or water quality. Given their strong dislike of biting insects and belief that Bti is the most natural and effective tool available, this perspective has been labeled *the Bti enthusiast*.

### 3.4. Convergence and divergence between all three social perspectives

While perspectives 1, 2, and 3 share certain viewpoints, they also reveal significant divergences in their attitudes toward the use of Bti and its implications. Perspectives 1 and 2 are the most closely aligned in terms of environmental and public awareness, as the correlation coefficients demonstrate (Table 3), both advocating for greater consideration of public awareness and alternative methods to Bti. However, only perspectives 1 and 3 seem to be aware of and convinced by the negative ecological impacts of Bti, particularly regarding its effects on the food chain. Nevertheless, for perspective 3, these environmental concerns do not outweigh the importance of human comfort, justifying the use of Bti. Interestingly, perspectives 2 and 3 appear more accepting of Bti, especially when its application is rigorously controlled. These two perspectives also converge on the notion that Bti has proven to be a more effective solution than repellents or protective clothing for improving human comfort and wetland appreciation. In contrast, perspective 1 places greater emphasis on the ecological role of biting insects, viewing their presence as essential to the functioning of aquatic ecosystems. As such, perspectives 1 and 3 represent the most polarized views, having the lowest correlation coefficient (Table 3), one prioritizing environmental integrity and the other human comfort. Perspective 2 occupies a more moderate position, expressing skepticism about the larvicide’s environmental risks, all while rejecting the idea that Bti could significantly harm pollinators, birds, or insectivorous species. Notably, all three perspectives express uncertainty or lack of knowledge regarding the potential impacts of Bti on human health and its toxicity. They express, some more than others, the need to share this knowledge with the population before spreading.

**Table 3.**
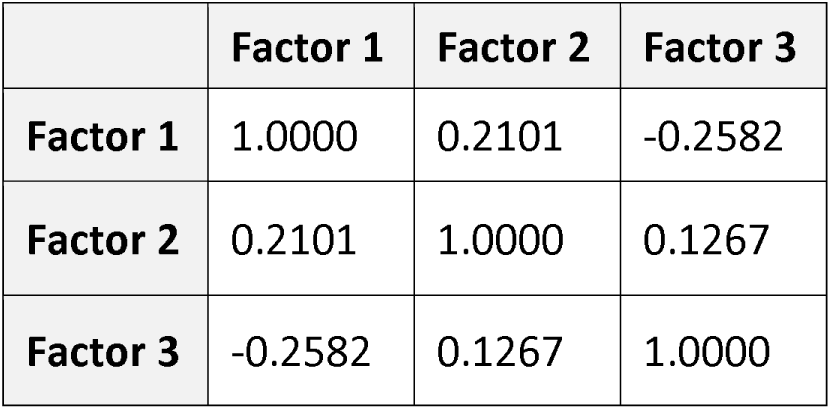
Factor correlation matrix. Factors correspond to each perspective, respectively.

## 4. Discussion

Our study provides the first in-depth exploration of key social discourses surrounding the use of Bti in urban regions of Quebec, filling a significant knowledge gap regarding the social dimensions of larvicide use in temperate and boreal regions. Using the Q methodology, we identified three distinct social perspectives on Bti-based mosquito and black fly control, advancing our understanding of public perceptions in environmental interventions, which remains underexplored, mainly in temperate climates. Unlike tropical and subtropical regions, where social aspects of vector control have been more widely studied, temperate and boreal regions have seen limited integration of social science into environmental management. This study also highlights the critical need for transparent communication, public awareness, and precautionary approaches in guiding future policies, thus providing a valuable foundation for more socially acceptable and ecologically responsible biting insect management strategies. Overall, this study contributes to the growing body of literature on pesticide use for biting insect control, particularly for Bti in regions where the health risks posed by biting insects are minimal.

Our study aimed to highlight the social perspectives that can shed light on why the general public accepts or rejects the use of Bti, given that it represents a low risk to human health but can have significant consequences for natural ecosystems. Our results demonstrated that the use of Bti to control mosquitoes and black flies in wetlands and water environments does not reach a consensus among the population. The reasons why respondents have diverging opinions on the use of Bti are diverse and related to personal values and norms; some individuals prioritize human comfort, particularly for enjoying outdoor activities, while others express environmental concerns or criticize the lack of public information. However, people agreed on the general lack of knowledge and certainty regarding the repercussions of using Bti in aquatic ecosystems.

Interestingly, most participants had no *a priori* knowledge of the current use or the nature of this larvicide. Their views were shaped primarily by the limited information provided by the team during the interviews. This widespread lack of public awareness highlights a critical issue that can lead to uninformed decision-making and the adoption of policies that may not reflect public values or ecological considerations. These results align with another Canadian study that revealed the lack of knowledge among the public and elected officials about Bti and its effects (Peterson et al., 2025). In Perterson et al. (2025), public opinion was divided between two perspectives: the fear of the environmental effects of Bti and the positive effects of Bti on human comfort and quality of life.

### 4.1. Social acceptability of the use of Bti

Understanding which factors influence the social acceptability of Bti is essential for implementing effective biting insect control programs while ensuring community support and ecological protection. Generally, the factors that have the most influence on social acceptability include the real or perceived risks, benefits and impacts on local communities, involvement in the decision-making processes, the level of trust in promoters and institutions, and broader social, economic, geographical, and territorial contexts (Ministry of Natural Resources and Forests, 2025.; Tanguay et al. 2021). Additional key elements include local knowledge, uncertainty, and the values, beliefs, and expectations of the population (Ministry of Natural Resources and Forests, 2025).

In our study, participants exhibited varying levels of social acceptability toward BtI use, even though there is a very low health risk. With clear and significant health hazards related to diseases transmitted by biting insects, human populations in warmer and tropical climates tend to have higher social acceptability towards Bti use. For example, in rural Burkina Faso, a qualitative study revealed a positive community perception and acceptance of Bti (Dambach et al., 2018). However, most participants in this study were unaware of the specific product used, which highlights the lack of public awareness. For most participants, the program was generally perceived as effective and successful in controlling malaria and reducing mosquito nuisance, facilitating routine implementation. In the Burkina Faso case study, respondents expressed their continued support for larvicide activities with a high willingness to financially contribute (Dambach et al., 2018). Similar results were shown in several other studies conducted in Tanzania (Mboera et al., 2014) and Rwanda (Munyakanage et al., 2025; Ingabire et al., 2017). While many citizens appreciate the reduction in biting insects, others, particularly in sensitive environments or near protected areas, express their opposition to a form of interference with natural ecosystems and would rather use alternatives (Brühl et al., 2020; Reuss et al., 2020). In temperate regions, social acceptability is likely to reflect the tolerance threshold for biting insects rather than the perceived risks of being bitten. Tolerance is based on individual perception, which is difficult to measure and anticipate, especially within different communities and cultures. While larvicides such as Bti may be justified in regions where vector-borne diseases pose significant public health risks, their widespread application in temperate and boreal regions, where the primary concern is human discomfort, may require greater scrutiny. The risks must therefore be put into perspective in a North American context.

### 4.2. Biodiversity crisis and public awareness

The global biodiversity crisis calls for rigorous assessments of the ecological impacts of larvicides and pesticides, alongside a greater consideration of eco-friendly and sustainable alternatives. Insect biodiversity loss, in particular, has long been underestimated and remains critical yet often overlooked, partly due to the vast diversity of insect species, as well as methodological challenges in obtaining representative data (Saunders et al., 2025). Nevertheless, growing evidence confirms that insect populations are declining globally, driven by multiple anthropogenic pressures including greenhouse gas emissions, agricultural intensification, habitat degradation, reduced plant diversity, landscape homogenization, and increased pesticide use (Saunders et al., 2025; John et al., 2024). Regrettably, insects have been marginalized in scientific research, conservation initiatives, and policymaking, partly due to their elusiveness, negative perception and low aesthetic appeal, which is often considered less charismatic than that of most vertebrates (John et al., 2024; Van der Sluijs, 2020; Troudet et al., 2017). The global biodiversity crisis is also contributing to ecological imbalances, notably through the loss of insect predators, like birds and amphibians, that help regulate insect populations and the disruption of feeding habits caused by climate-induced shifts in seasonal timing (John et al., 2024).

Alternatives such as natural insect repellents, nets, traps, protective products, and specialized clothing are commonly used to reduce bites and other irritations. While these approaches still demand a certain level of tolerance and patience due to their limited efficacy and duration, they can still contribute to reducing reliance on larvicides and insecticides. To help encourage these alternatives and improve tolerance, public awareness and education are key. It is essential to acknowledge the critical roles insects play in food security and public health, including the decomposition of organic matter, nutrient cycling, soil fertility, and crop pollination (John et al., 2024; Van der Sluijs, 2020). Insects play a vital role in both food production and the development of phytopharmaceuticals and nutritional supplements, as well as in other key ecosystem functions (John et al., 2024; Van der Sluijs, 2020). That being said, public awareness and increased concern for the biodiversity crisis will likely help promote alternatives and guide environmental interventions, such as Bti application in aquatic ecosystems.

### 4.3. Adopting systems thinking and the precautionary principle

In temperate and boreal regions, we should adopt a systems thinking approach regarding the use of Bti. Systems thinking is a concept that enables us to understand the complexity of ecosystems and social interactions, providing a more holistic perspective. It allows us to see beyond the direct effects of Bti. This means considering not only the desired effect of Bti, *i.e.,* the reduction of mosquitoes and black flies, but also its indirect consequences and cascading effects. By applying this concept, it is possible to assess the impacts on non-target species as well as the food web, such as a decrease in mosquito predators. This concept has already proven its worth in environmental policy. The study conducted by Phan et al. (2025) demonstrated how systems thinking has enabled the identification of various interactions that inform decision-making in aquatic ecosystem conservation. Indeed, in this case study, the various interactions between agricultural production, pesticide use, water quality improvement policies, and climate change were examined. Understanding these interactions provided insight into the structure and functioning of this complex system, identifying key points of influence for strategic decision-making, such as water quality improvement policies that can limit terrestrial pollutants from entering aquatic environments.

As for the precautionary principle, often invoked in environmental governance, this principle asserts that scientific uncertainty should not delay preventive action in the face of potential harm *(Rio Declaration 1992, Principle 15)*. The need for both systems thinking and the precautionary principle is especially evident when considering the disconnection between the implementation of Quebec’s *Wetland and Water Conservation Act* (Assemblée Nationale, 2017; RLRQ, c. Q-2, r. 9.1) and the spreading of larvicides in these water environments that provide essential ecosystem services. On the one hand, the governance claims to be committed to biodiversity protection goals (Ainsworth et al., 2022), and on the other, it sometimes promotes the application of Bti programs in highly diverse aquatic habitats. These decisions pose a contradictory statement, as studies demonstrate both direct and indirect impacts of Bti on the environment and unintended species. By combining systems thinking and the precautionary principle, it is possible to overcome the inconsistency in public actions and better manage uncertainties and gaps resulting from a lack of knowledge. This approach promotes the integration of sustainable management by aligning short-term decisions with long-term ecological and social objectives. Biting insect control programs should not be treated as an isolated solution, but rather as an intervention applied in a complex socio-ecological system. It is therefore imperative to implement strategies that do not undermine the targeted conservation objectives.

### 4.4 Study limitations

This study was conducted primarily in an academic context by undergraduate ecology students using a standardized and structured methodology, including a fixed interview script to reduce variability. However, certain limitations must be acknowledged. Many participants were recruited from the students’ networks, such as family and friends, which may have introduced selection bias. This could have led to an overrepresentation of individuals with pro-environmental values, potentially influenced by social desirability bias or pre-existing alignment with the interviewers’ perspectives. Social desirability bias refers to the tendency of respondents to provide answers they perceive as socially acceptable, rather than responses that accurately reflect their true thoughts or feelings, in order to avoid judgment from others (Podsakoff et al., 2003). This bias is recognized as a common source of measurement error in behavioral research, especially when sensitive or normative issues are addressed (Podsakoff et al., 2003). These factors may partly explain the higher representation of the first perspective, “The Critical Environmentalist.” While the study offers valuable insights, future research should aim to include a larger and more randomized sample of citizens to enhance the representativeness of a city-wide population.

## 5. Conclusion

The use of Bti as a tool to control biting insects in Quebec illustrates a classic tension between public health, human comfort, and ecosystem preservation. While Bti offers an apparently low-impact and targeted solution, its long-term effects on aquatic biodiversity remain unclear. From a sustainable development perspective, it seems essential to adopt a precautionary, adaptive, and transparent approach, based on strong scientific evidence, and to integrate complementary alternatives: landscaping, individual protection, and information campaigns. As social perspectives continue to shape environmental governance, it becomes essential to integrate both ecological evidence and societal values into decision-making that will support more sustainable and precautionary policies. Taking these social perspectives into account can provide valuable insights for the current debate on Bti use in urban areas in Quebec, and elsewhere, and inform future decision-making processes.

## Annexe

**Appendix A.** List of the 29 themes identified during the group thematization exercise from the thematic analysis.

| Justifications against Bti (n=15) | Justifications in favor Bti (n=13) |
| --- | --- |
| Alternatives ot Bti exists | Improve quality of life |
| Existential reasons (e.g. mosquitoes and flies have the right to exist) | Insects are invasive |
| Risk of developing resistance | Mosquitoes are a nuisance |
| Bti is a chemical product | Control public health issues |
| Mosquito season doesn't last long | Necessary in targeted areas |
| Possible impact on Bti on water quality | An alternative to chemical pesticides |
| Tolerance to mosquitoes must be developed | With greater control and stricter regulations |
| Mosquitoes are not harmful | Mosquitoes are a source of dirt for homes |
| Against application of Bti, but may be considered depending on situation | Less damaging to the environment |
| Bti is too expensive | Does not affect the environment |
| Bti is not very effective | Possibility for wildlife of finding other food |
| Studies on Bti are insufficient | Doubts about Bti effectiveness |
| Importance of maintaining natural cycle | Improve appreciation of wetlands |
| Accumulation in the environment |  |
| Not good for biodiversity |  |

**Appendix B.**
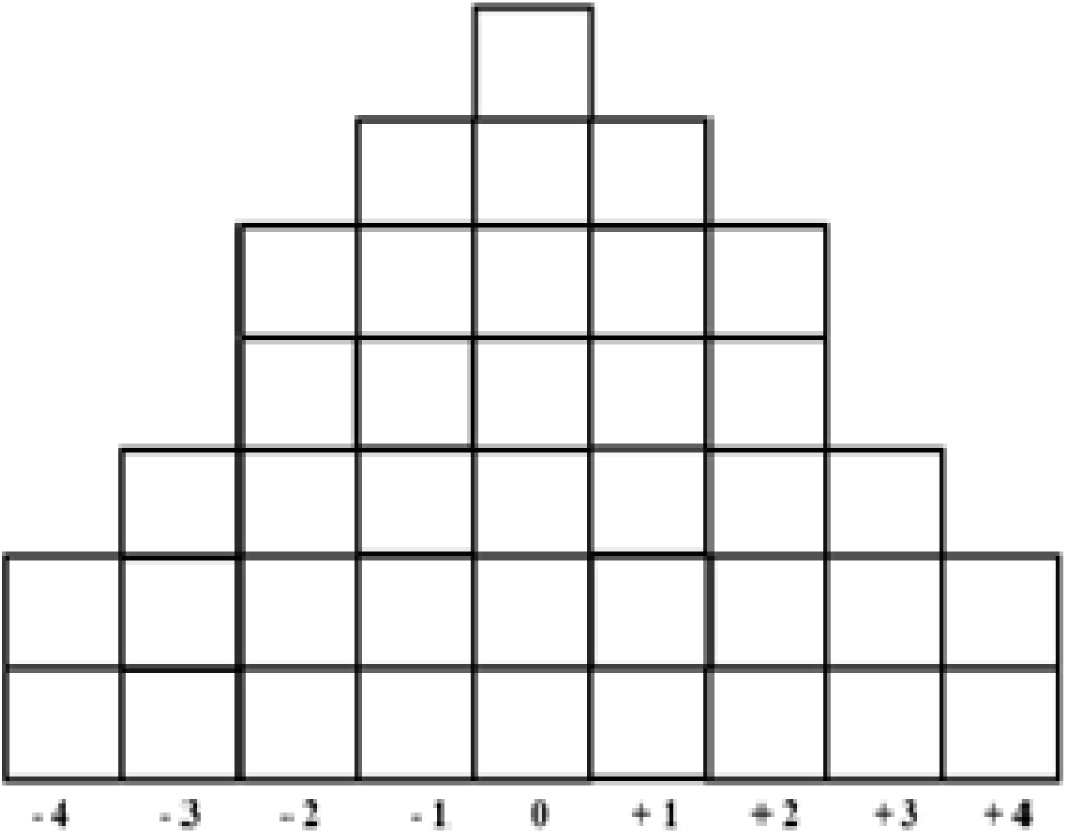
Q-sets: pyramid-shaped rankings of statements based on participants’ levels of agreement.

**Appendix C.**
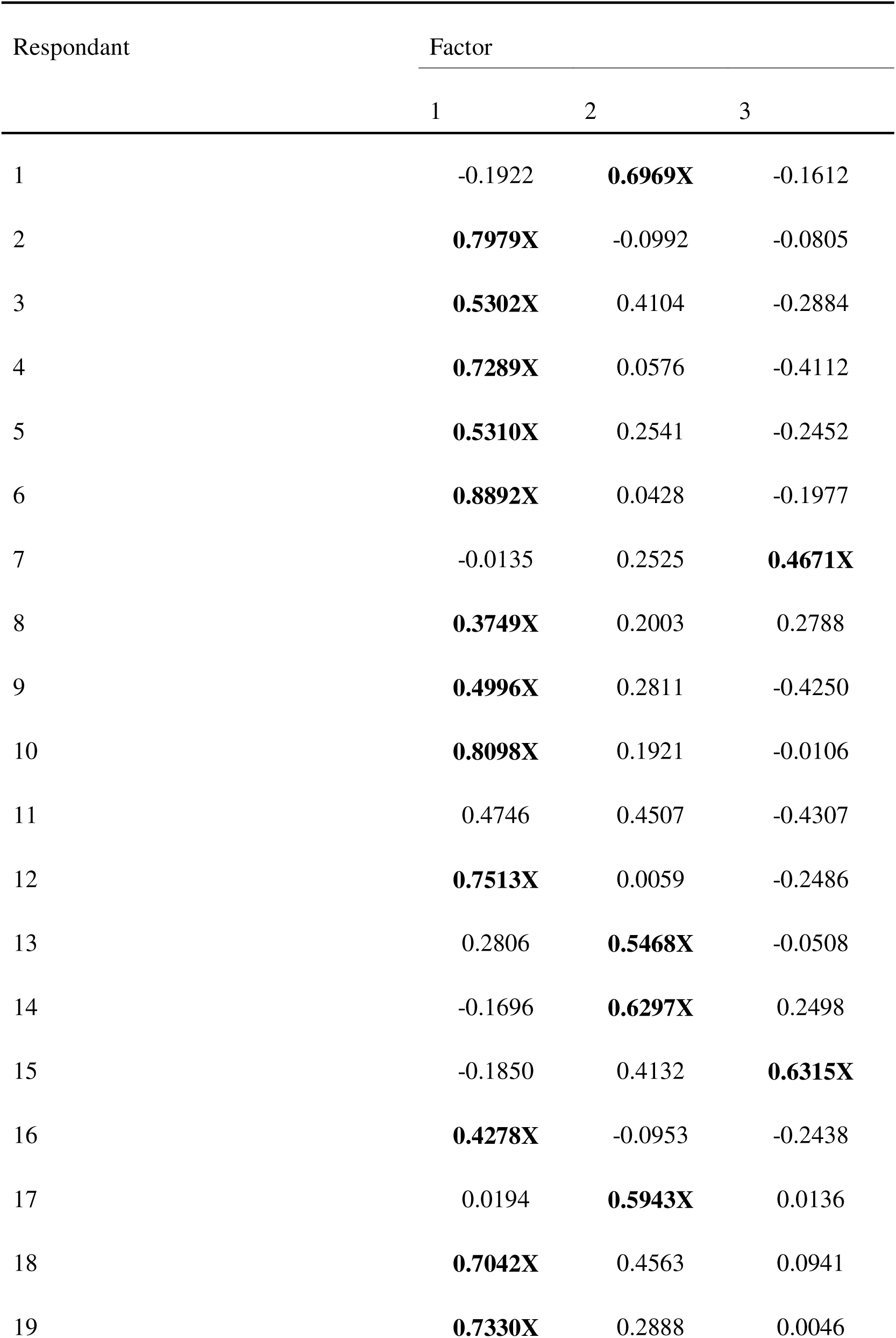
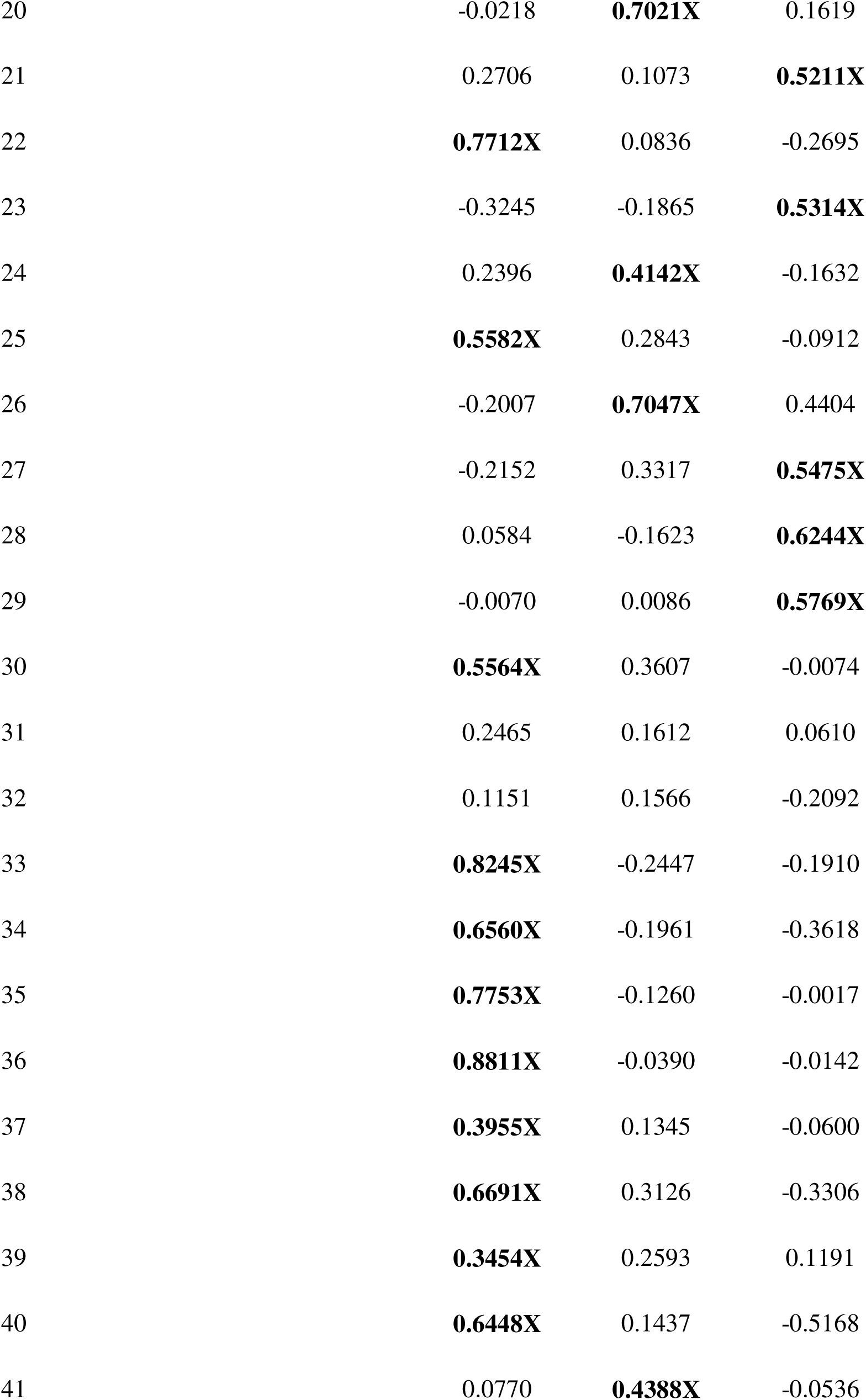
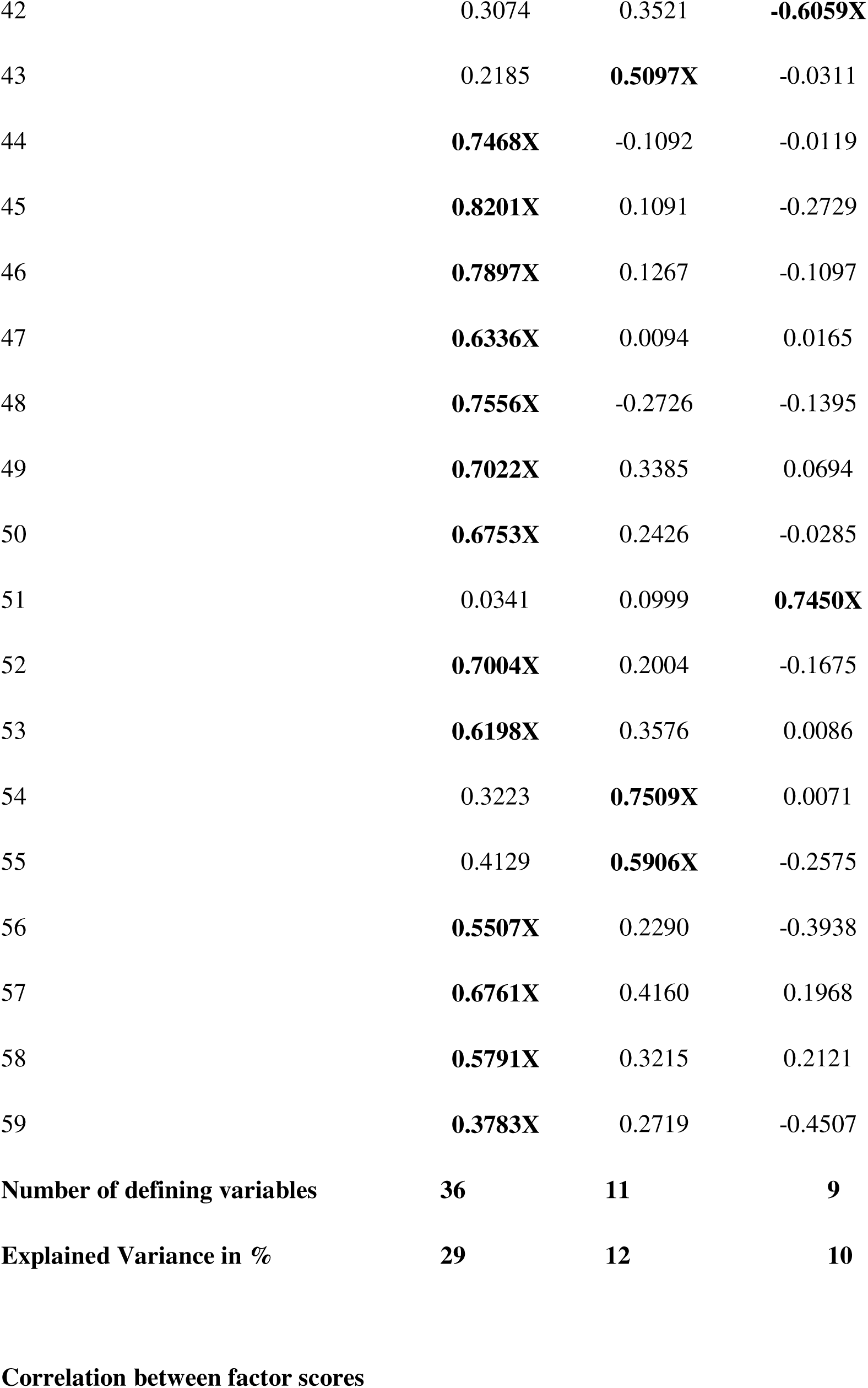
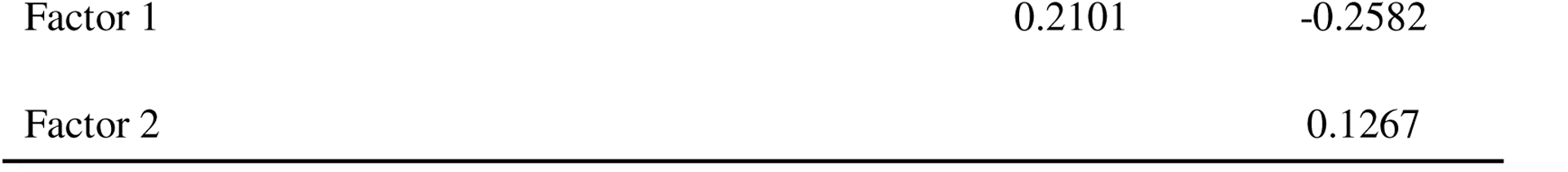
Respondent profiles corresponding factor scores. Scores in bold and marked with an X are those used to estimate the factors.

**Appendix D.**
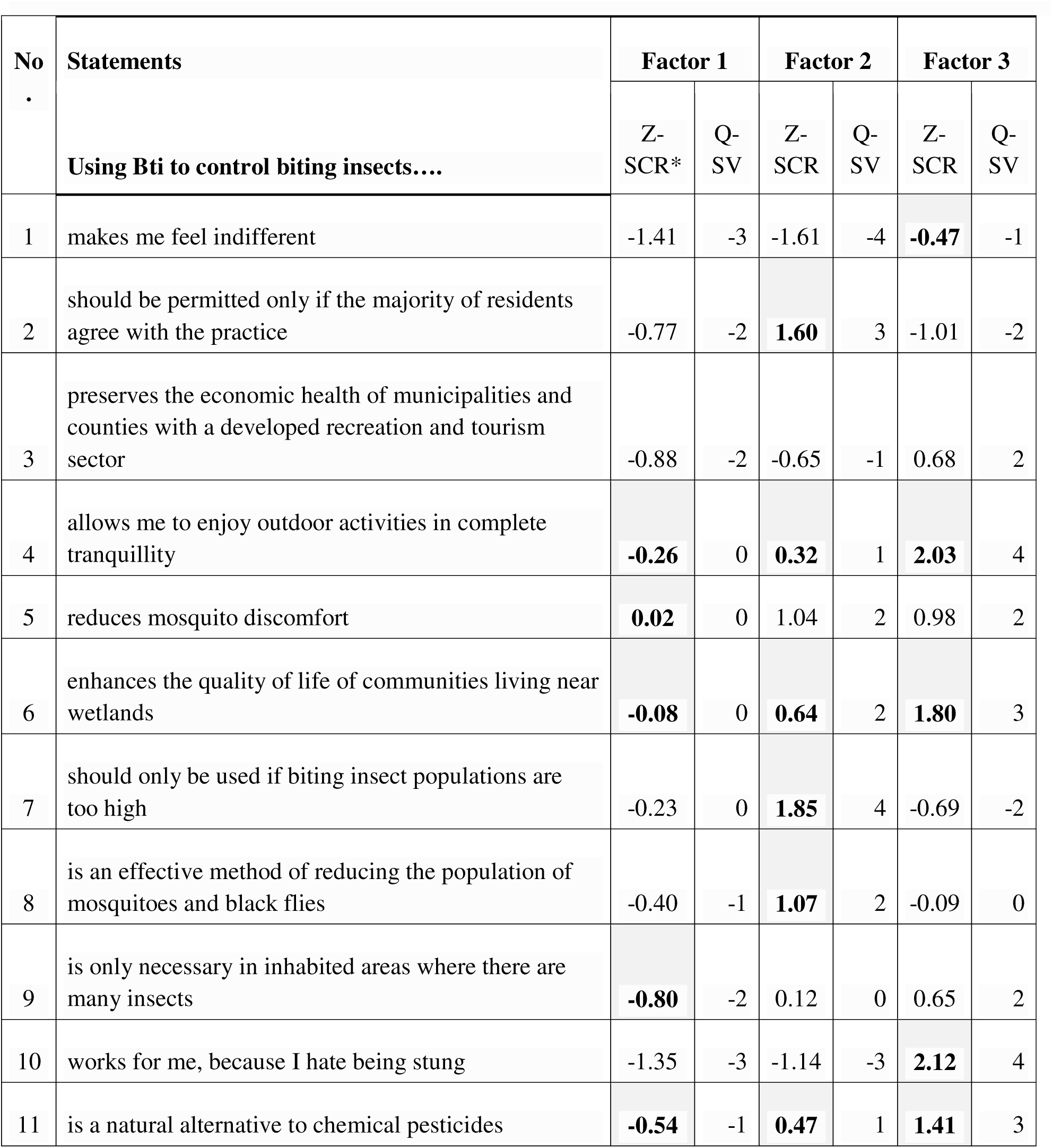
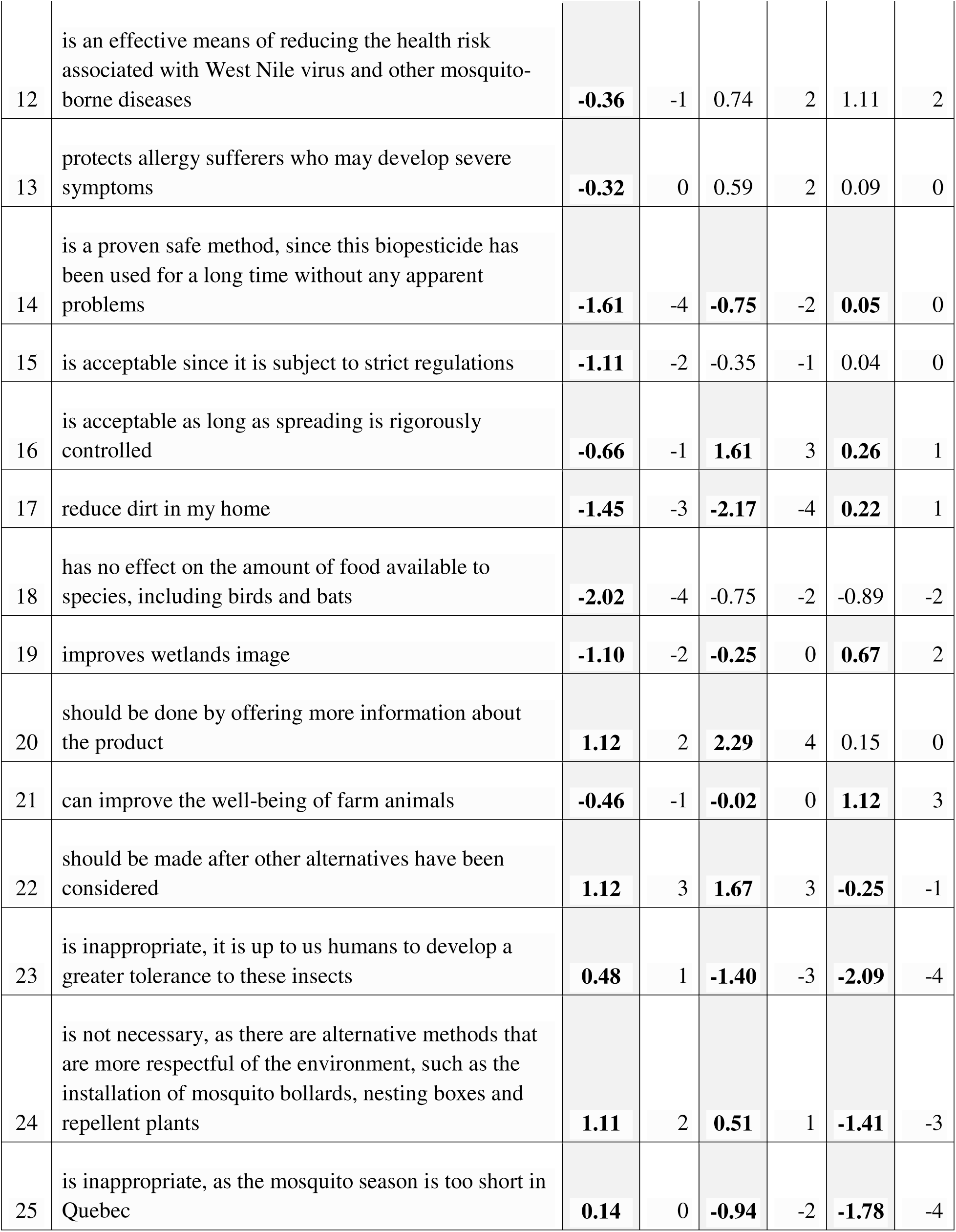
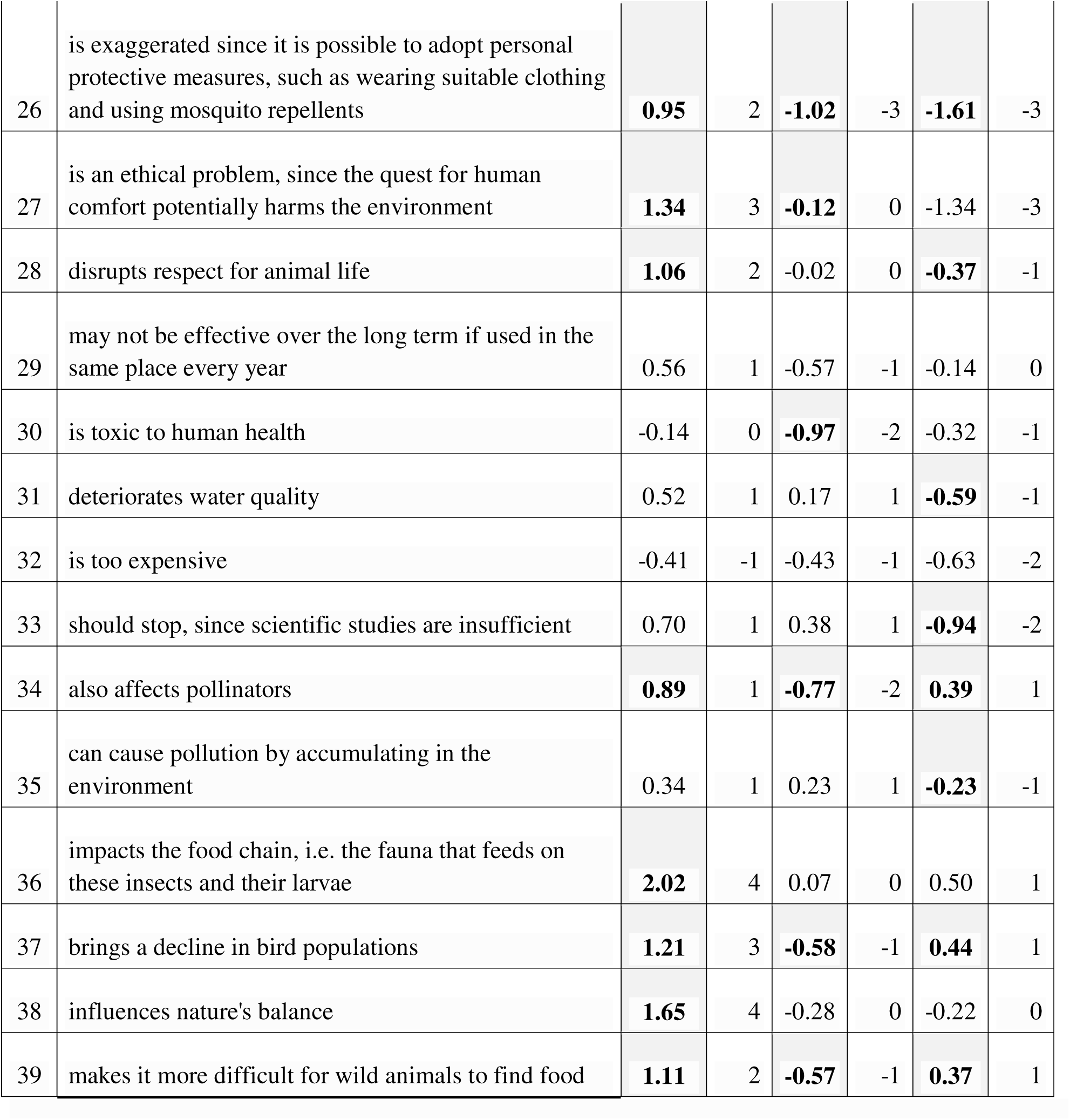
Factor z-score of statements with ranking. Z-scores in bold and gray are the distinctive statements for each factor. *Z-scores are standardized scores (Watts & Stenner, 2005). It allowed us to determine the relationship between the statements and factors.

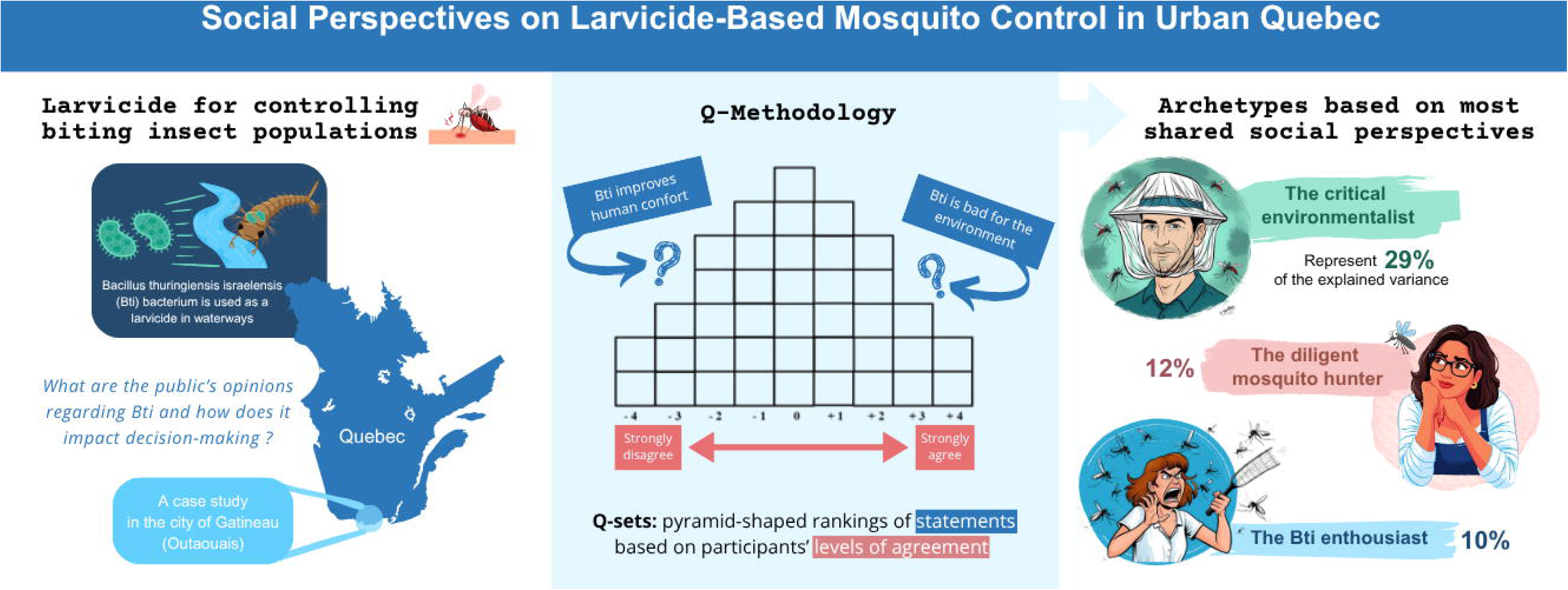

